# An agent-based model of adaptation of holobionts with different microbial symbiont transmission modes

**DOI:** 10.1101/2020.10.08.330902

**Authors:** Siao Ye, Zhu Liu, Evan Siemann

## Abstract

The hologenome theory suggests that holobionts (host plus symbiont) with hosts that are only able to adapt slowly may be able to persist in deteriorating environmental conditions via rapid adaptation of their microbial symbionts. The effectiveness of such symbiont adaptation may vary depending on whether symbionts are passed directly to offspring (vertical transmission) or acquired from the environment (horizontal transmission). However, it has been suggested that holobionts with horizontal transmission cannot pass down their symbionts faithfully, preventing adaptation at the holobiont level because of host-symbiont disassociation between generations. Here we used an agent-based model to investigate whether holobionts with horizontal microbial symbiont transmission can adapt to increasing stress solely through symbiont adaptation and compared their adaptation to holobionts with vertical transmission. We found that holobionts with either transmission mode were able to adapt to increasing abiotic stress solely via symbiont adaptation. Moreover, those with horizontal transmission were more competitive than those with vertical transmission when hosts were able to selectively associate with the most suitable symbionts. However, those with horizontal transmission were less competitive than those with vertical transmission when symbiont establishment was random. Our results support the hologenome theory and demonstrate that holobionts with horizontal microbial symbiont transmission could adapt to increasing abiotic stress via their symbionts. We also showed that whether holobionts with horizontal or vertical symbiont transmission are favored in increasingly stressful conditions depends on the ability of hosts to recognize and foster microbial symbionts that confer stress tolerance.

**IMPORTANCE:** Symbiotic organisms such as reef building corals are sensitive to environmental perturbations due to anthropogenic disturbances or climate change, and it is critical to understand whether they are able to adapt to previously unfavorable conditions. To date, studies have focused on the impacts of existing microbial symbiont variation on holobiont stress tolerance but here we use agent-based models to explore holobiont adaptation via symbiont adaptation. We studied both deterministic and stochastic processes in holobiont adaptation by investigating the following four factors: holobiont transmission modes, a host’s ability to recognize tolerance-conferring symbionts, a symbiont’s mutational variance, and rate of stress increase. Our simulation provides a comprehensive understanding of holobiont adaptation under stress, which not only has implications for future endangered symbiotic species management, but also provides fresh insight into species evolution as proposed by the hologenome theory.

## INTRODUCTION

Research on symbiotic organisms has suggested that symbiont variation can drive holobiont (host plus symbiont) phenotypic diversification, and influence holobiont fitness (1). In addition to host genetic changes, symbiont genetic changes can alter the holobiont phenotype and therefore impact holobiont adaptation (2). Zilber-Rosenberg and Rosenberg proposed that rapid holobiont adaptation could take place even when host adaptation is slow, because symbiont adaptation and composition shifts could be rapid (3). A diversity of potential symbiont partners are rich resources for holobionts, which might be critical for holobionts to adapt to increasing environmental stress such as global warming (4, 5).

Compared to hosts, symbionts have large population sizes and short generation times, and may adapt to ambient changes (6), yet it is unclear whether symbiont adaptation alone could drive holobiont adaptation. Most studies on holobionts focused on variation in symbiont composition rather than symbiont adaptation. For example, researchers found cnidarians associating with different symbiont strains differ in their thermal tolerances (5, 7), and thermal tolerance of aphids can be controlled by their symbiotic bacteria (8, 9). Such changes in holobiont phenotype are mainly based on existing symbiont diversity and can be observed within one generation. However, experiments testing holobiont adaptation via symbiont adaptation and composition shifts would be difficult because they require tracking of symbiont composition as well as holobiont phenotypes over a long time and so is less explored. Chakravarti et al. (10) pointed out that assisted evolution in symbionts can be applied to improve holobiont stress tolerance, and they were able to detect its impact on symbionts stress tolerance, but not on the holobiont stress tolerance. Some field observations also suggest holobiont adaptation can be achieved through symbiont adaptation. Rodriguez et al. (11) discovered the stress tolerance conferred by a symbiotic fungi to host plants is habitat dependent. Another study on corals also revealed local adaptation existed in symbionts and leads to holobiont divergence in their thermal tolerance, in which the symbiotic algae collected from warmer reefs maintained higher resistance to heat compared to those from cooler habitats even after multiple asexual generations (12). Nevertheless, it is hard to monitor symbiont adaptation in the field over evolutionary time scales and track its impact on holobiont fitness. A simulation model would be appropriate to study holobiont adaptation via symbiont adaptation (13, 14), which is lacking in holobiont research.

To investigate whether holobionts can adapt to stressful environments via symbiont adaptation, it is critical to understand whether holobionts with either horizontal symbiont transmission (“H” hereafter) or vertical symbiont transmission (“V” hereafter) are both able to pass their adaptive symbionts to offspring, and which type is better at coping with stress. Some researchers argue that passing down adaptive symbionts to offspring is unstable for holobionts conducting horizontal transmission, which makes the hologenome theory less plausible (15, 16). They suggest the symbiont community could be shaped simply by opportunity and environmental filters, and the transmission fidelity is low (15, 16). In addition, opposite opinions are given when discussing H’s and V’s adaptability. In vertical transmission, individuals inherit symbionts directly from parents, which is similar to classical gene inheritance, so adaptive symbionts from parents can be passed to offspring (17). But there are drawbacks associated with this transmission mode; symbionts picked up by offspring might only be a subset of their parents’, so symbiont diversity could decrease dramatically through evolutionary time, and become maladaptive once the ambient conditions change (18, 19). On the other hand, individuals conducting horizontal transmission pick up a variety of symbionts in the neighborhood, so they might respond to environmental changes faster by associating with stress tolerant symbionts (4, 20). However, the disassociation of symbionts and hosts between generations could lead to loss of adaptive symbionts and colonization of virulent symbionts (21, 22). Most studies on microbe transmission and adaptation in the past investigated how different transmission modes affect virulence (17, 22–24), and explored the dynamics of virulence-transmission trade off instead of considering them as a selection unit (25–27), because their systems emphasize antagonistic rather than mutualistic relations between the host and the symbiont. Roughgarden (28) conducted a pioneering simulation of holobiont evolution and found both vertically transmitted and horizontally transmitted agents were able to evolve as a holobiont unit. But she used the number of microbes to determine holobiont fitness in her model, and did not study the interactions between the environment and holobionts. Because of the complex interactions among the host, the symbiont, and the environment, combined with the different symbiont acquisition and retention mechanisms in Hs and Vs, it is hard to predict whether Hs and Vs are able to adapt to climate change, and which mode is more adaptive with increasing environmental stress. Holobionts with different transmission modes could differ in their novel symbiont acquisition and retention, which makes it hard to predict whether they are able to adapt to climate change, and which modes is more adaptive.

Here we use agent-based modeling to explore how holobionts with different transmission modes respond to increasing stress because such models are flexible and able to handle complex problems (29, 30). Interestingly, such topics are better explored in cultural transmission, where researchers explore how knowledge information (*e.g*., language, knowledge, etc.) is transmitted between generations under different environment regimes (31). In cultural transmission, vertically inherited information (from parents) is more conservative than obliquely acquired (from elder people excluding parents or ancestors) (32), which resembles vertically transmitted symbionts and horizontally transmitted symbionts. Some studies reveal oblique transmission is favored when the environment fluctuates, and is necessary in driving language evolution, while vertical transmission is preferred when the environment is stable (33, 34). However, there could be fundamental differences between cultural transmission and symbiont transmission. For example, there is no increasing stress such as global warming in cultural transmission, so these studies do not provide insights into whether both transmission modes allow holobiont adaption to directional selection. In addition to transmission modes, we considered three other important factors that might impact holobiont persistence in changing climate: 1) the host’s ability to select adaptive symbionts, 2) the rate at which stress increases, and 3) symbiont mutational variance.

We propose that the ability to select adaptive symbionts is a key factor when studying holobiont adaptation, even though it is barely discussed, if at all, in previous research. Because the amount of resources and space an organism can provide to symbionts is limited, holobiont fitness is largely determined by the relative amount of adaptive symbionts within it (3, 35). So if symbiont acquisition is completely random, then Hs will have no advantages over Vs even if Hs have access to a larger symbiont pool from which they can choose partners, because the chances of getting adaptive and maladaptive symbionts are the same. In addition, drift might have larger impacts on Vs because the bottleneck effect is stronger in them due to the smaller effective symbiont population size (18). On the contrary, if symbiont acquisition is not random but can be determined by the host, then Hs might be more likely to associate with adaptive symbionts than Vs, and drift impacts would be limited. Such symbiont selection by hosts is not uncommon and has be found in both Hs and Vs, for example, solitary wasps block transmission of nonnative symbionts to offspring, and sorghum selectively increases monoderm bacteria during drought to improve plant growth (36–38). Thus we tested the impacts of drift on Hs and Vs adaptation by switching on and off the ability of holobionts to select for adaptive symbionts in our model.

Whether holobionts are able to adapt to the changing environment via symbiont adaptation also depends on how fast stress increases and how fast symbionts adapt. Holobionts such as corals are assumed to rely on the symbiont rather than the host to adapt to rising sea temperature, because corals are living close to their thermal limits and the temperature is increasing quickly (39, 40). Compared to the host, the symbionts are abundant with short generations, so they are more likely the key to holobiont adaptation (3, 41). Because most symbionts (*e.g*., bacteria, algae, fungi) reproduce asexually (10, 42), we consider mutation the major source of generating novel traits. Incorporation of mutation in our model makes the traits continuous and dynamic instead of static and discrete, and generates intra-individual as well as inter-individual symbiont variation. Such variation is necessary in understanding the complex interactions between holobionts and the environment, and interactions among holobionts (43, 44). Because population trait variance is sensitive to the magnitude of mutation (*i.e*., mutational variance) rather than mutation rate, it is important to know how mutational variance might affect distribution of symbiont traits and thus holobiont adaptation (45, 46). Large mutational variance may have greater impacts on horizontally transmitted holobionts, because once an extreme symbiont strain arises, it can be transmitted throughout the population and may be hard to lose, while it will be constrained in certain lineages in vertically transmitted holobionts. But large mutational variance could also produce lineages that have accumulated extreme symbionts in Vs when drift is strong. By varying the speed at which stress increases and mutational variance, combined with controlling drift, we explored how the selection-mutation-drift balance interacted with transmission modes.

Our goal was to present a model to test whether holobiont adaptation can be driven by associated symbiont changes, which involves symbiont mutation, acquisition, amplification and transmission. The agent-based model we used simulated changes at the environment level, the holobiont level, and the symbiont level, by varying four main parameters: transmission mode, the ability to select adaptive symbionts, the rate of environmental change and the magnitude of symbiont mutation. It involves interactions between the environment and the holobionts, among the holobionts, and between the holobionts and the symbionts. We believe our model could produce novel insights into holobiont adaptation.

## RESULTS

### Test case

We verified that when agents were selectively neutral in our basic model, the fixation was random at a 1:1 V to H initial ratio (Fig. 1).

**Figure 1:**
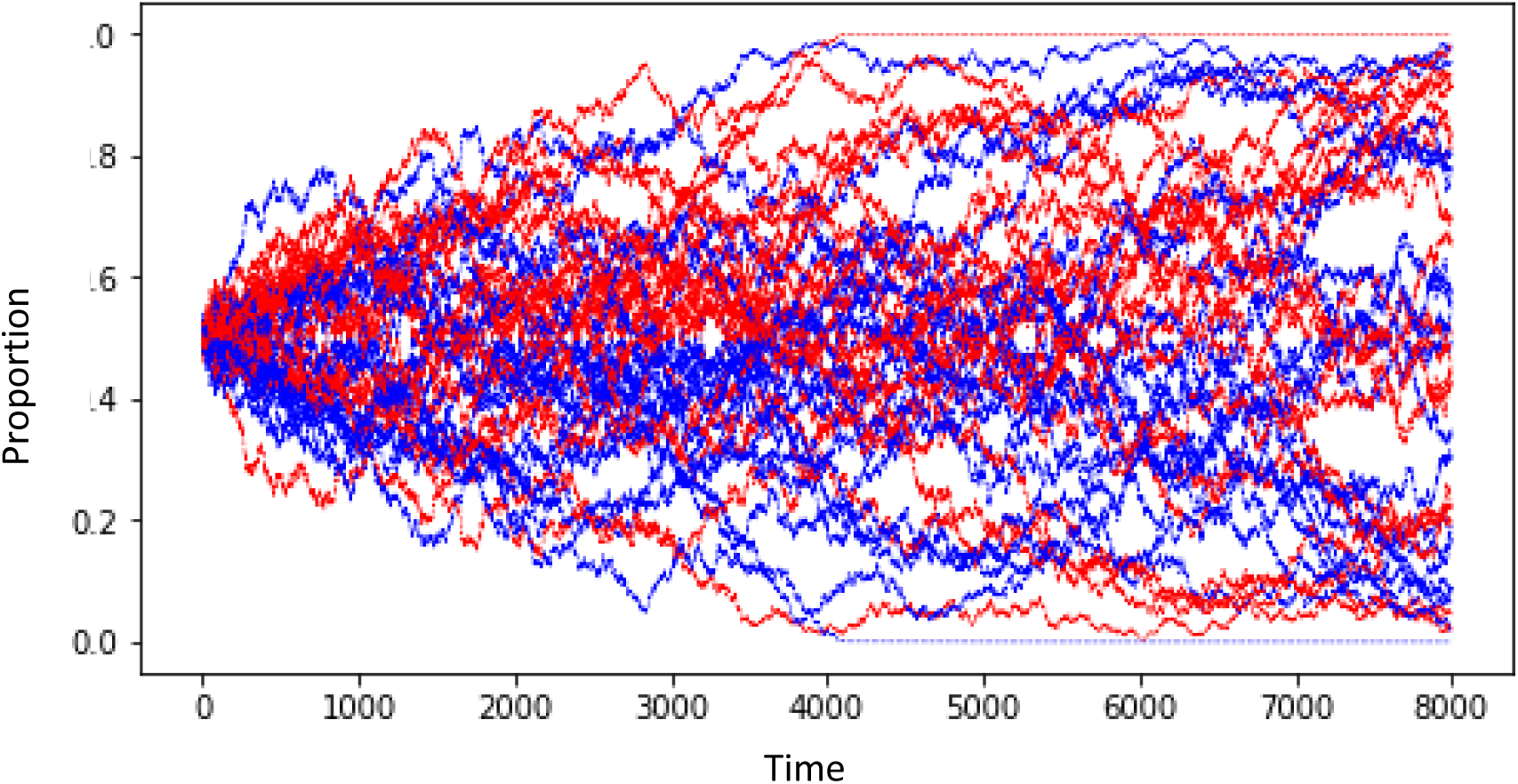
Fixation of holobiont transmission types under selectively neutral conditions and equal starting abundances of vertical and horizontal transmission types. Blue lines indicate the proportion of holobionts that have vertical transmission (V) and red lines indicate the proportion that have horizontal transmission (H). Each simulation is represented by a single blue line and single red line.

### Increasing temperature

Regardless of whether transmission is random or optimal, both Vs and Hs were able to adapt to increasing temperature when the rate of temperature increase was low and mutation variance was high (Fig. 2).

**Figure 2:**
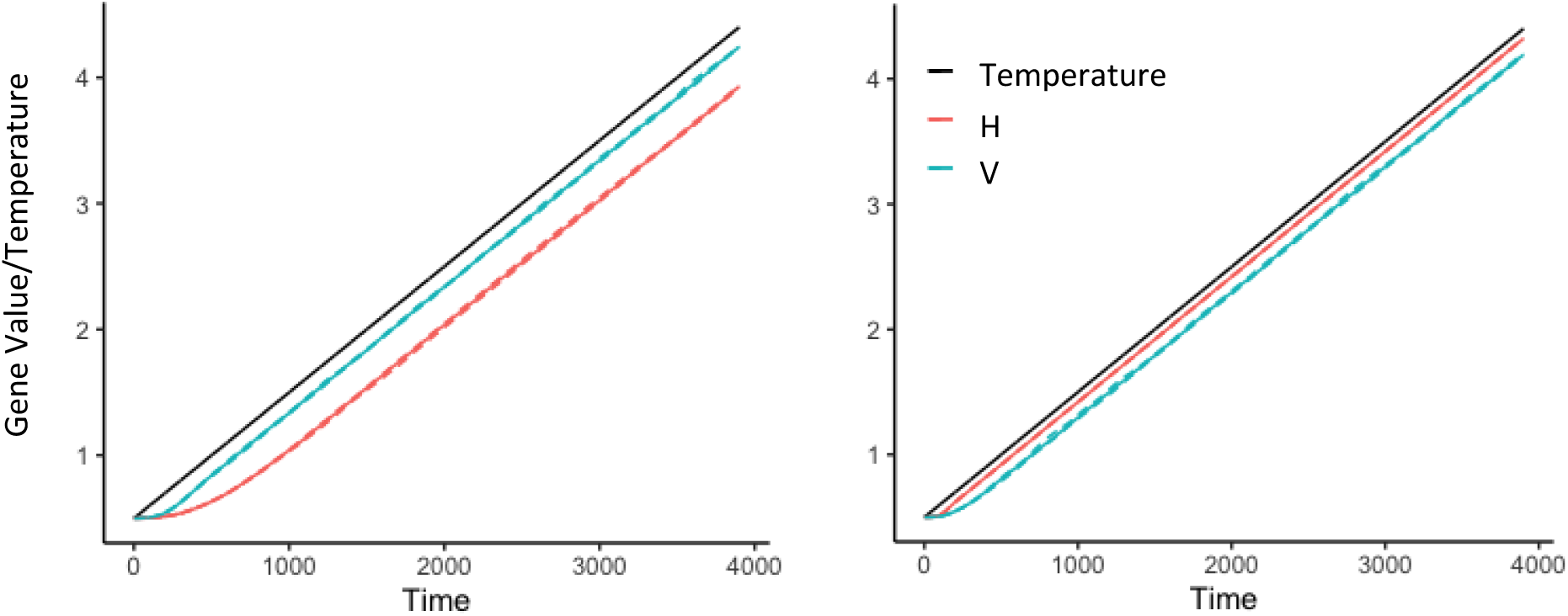
Average symbiont gene values (optimal temperature value) of horizontal (H) and vertical (V) transmission mode holobionts in (a) the random transmission model with temperature rate of change per time step=0.001 and magnitude of mutation with standard deviation=0.1, (b) the optimal transmission model at step=0.001, mutation_sd=0.01.

Generally, holobionts persisted in high mutation variance and low temperature rate of change conditions, but went extinct in low mutation variance and high temperature rate of change conditions.

However, the transmission mode did affect the relative fitness of Hs and Vs (Fig. 3). When transmission was completely random, Vs had higher fitness (Wilcoxon test, p=0.0002) and longer time to extinction (Wilcoxon test, p=0.03) compared to Hs for the same parameter combination, and their gene values also had higher correlation with the temperature (Wilcoxon test, p=0.0017). When transmission was optimal, Hs had higher fitness and longer time to extinction, as well as higher correlation between gene values and temperature.

**Figure 3:**
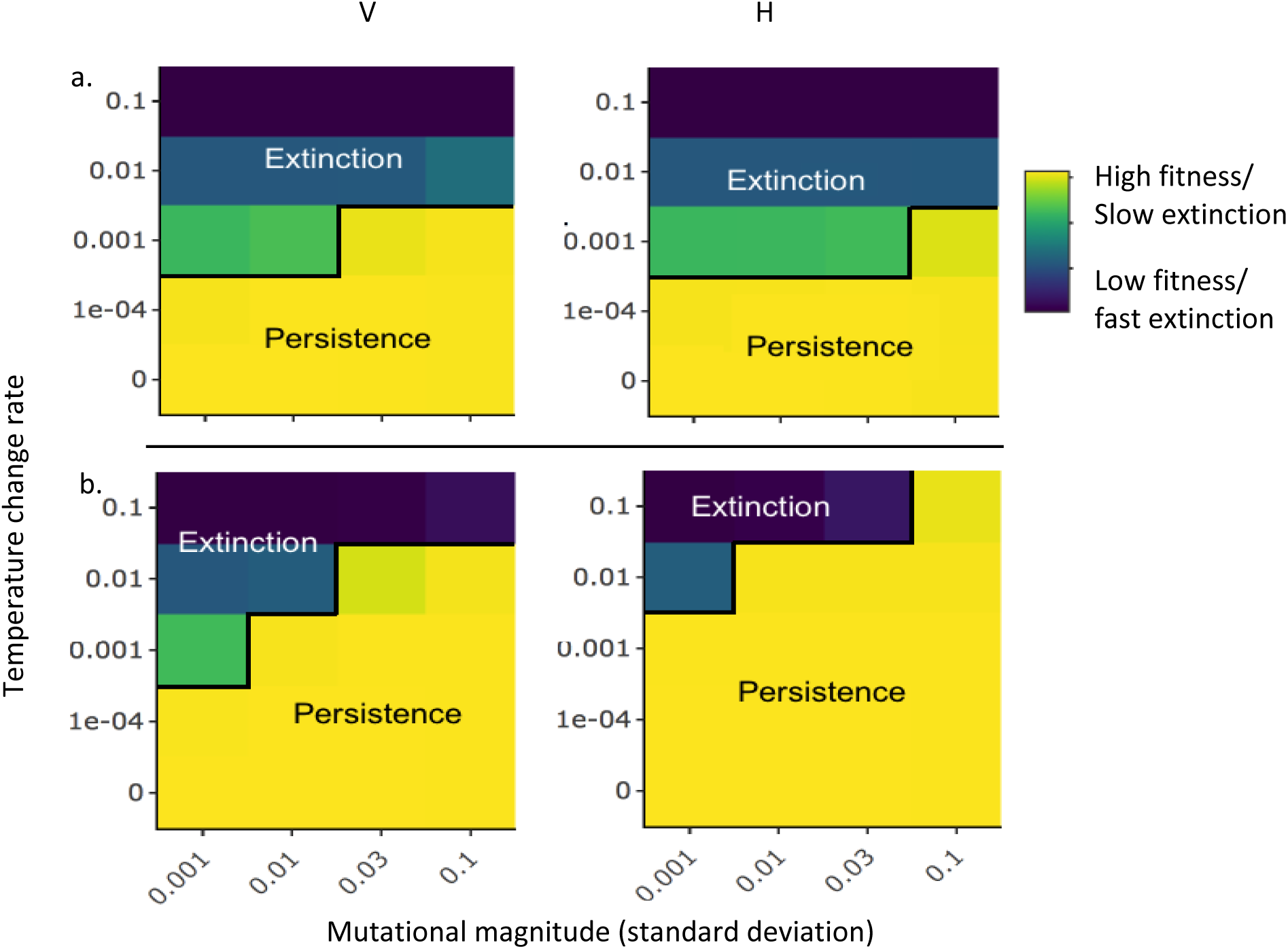
The dependence of fitness and time to extinction of vertical (V) or horizontal (H) transmission holobionts in separate simulations on rate of temperature change (magnitude of change per time step) and mutation magnitude (standard deviation) with (a) the random transmission model, (b) optimal transmission model. Rapid temperature change (large change per time step) and small magnitude mutations (low mutation variance) are likely to lower holobiont fitness and lead to extinction. The scenarios with persistence are those with high average fitness and those with low fitness have rapid extinction.

When both agent types were present at the same time, we were able to compare the ratio of Hs and Vs (Fig. 4), which tells their relative competitiveness. In both random transmission and optimal transmission models, Hs and Vs became extinct in large step/small mutation_sd regions, and coexisted in small step/small mutation_sd regions. In the rest of regions, Vs persisted in the random transmission model (Fig. 4a), and Hs persisted in the optimal transmission model (Fig. 4b).

**Figure 4.**
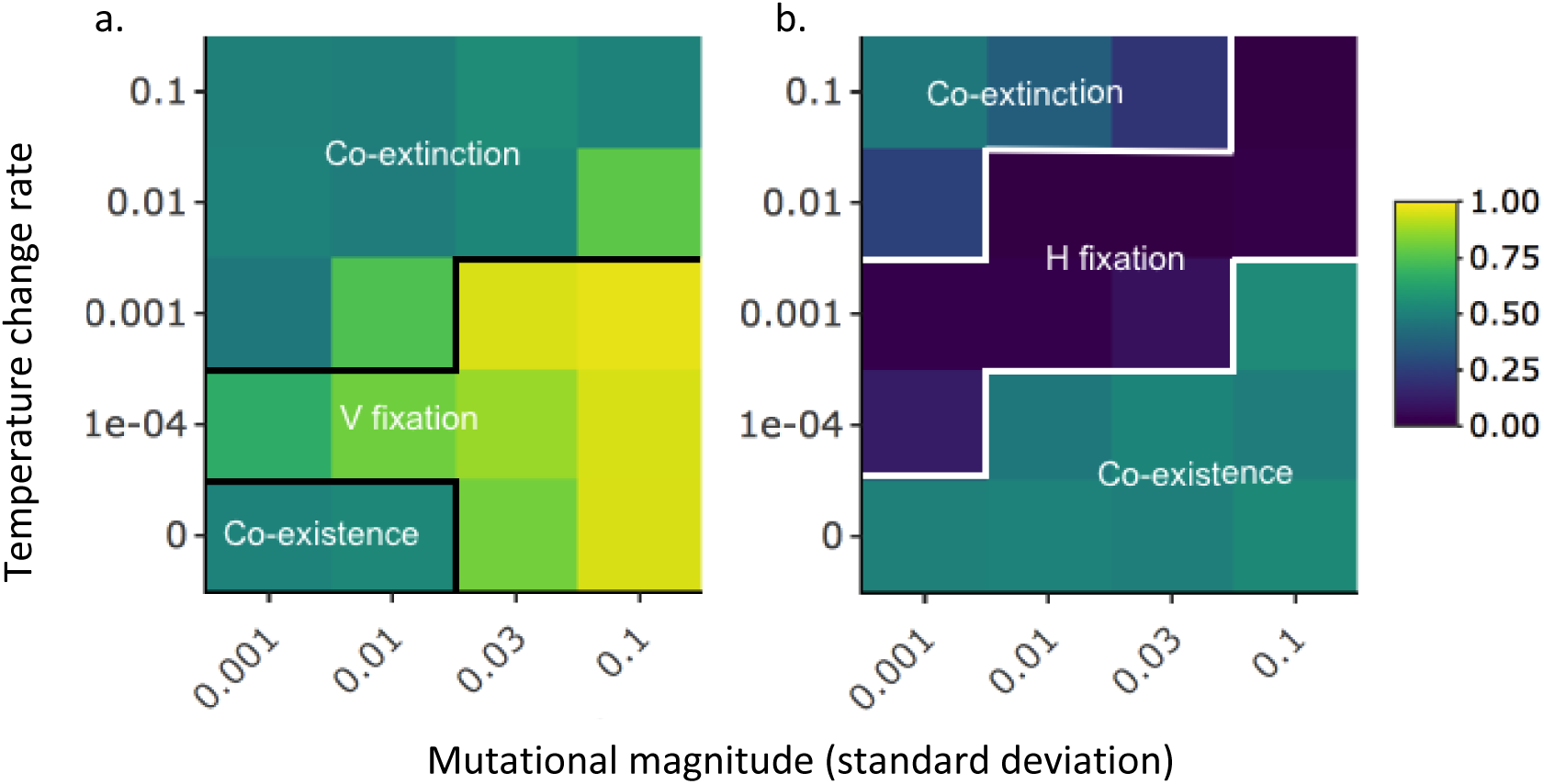
The dependence of average proportion of vertical transmission holobionts in simulations that include both vertical and horizontal transmission types on rate of temperature change (magnitude of change per time step) and mutation magnitude (standard deviation) with (a) random transmission, (b) optimal transmission. Brighter colors indicate conditions in which vertical transmission types reached fixation more quickly (and horizontal types became extinct) and darker colors indicate conditions in which vertical transmission types became extinct faster (and horizontal types reached fixation).

We also checked how virulence would affect the relative competitiveness of Hs and Vs by reducing birthrate of Hs by 5% at mutation_sd=0.001 and step=0 in both models. This resulted in Vs fixation in both models.

To explore how drift shapes symbiont distribution in the random transmission model, we compared fitness variance between individuals and within individuals (Fig. 5). We found Vs had larger variance among individuals (Wilcoxon test, p<0.001), but less variance within individuals (Wilcoxon test, p<0.001). This means symbiont distributions in each vertical holobiont were more homogenous than in each horizontal holobiont, but vertical holobionts were more heterogeneous than horizontal holobionts across the holobiont fitness landscape.

**Figure 5:**
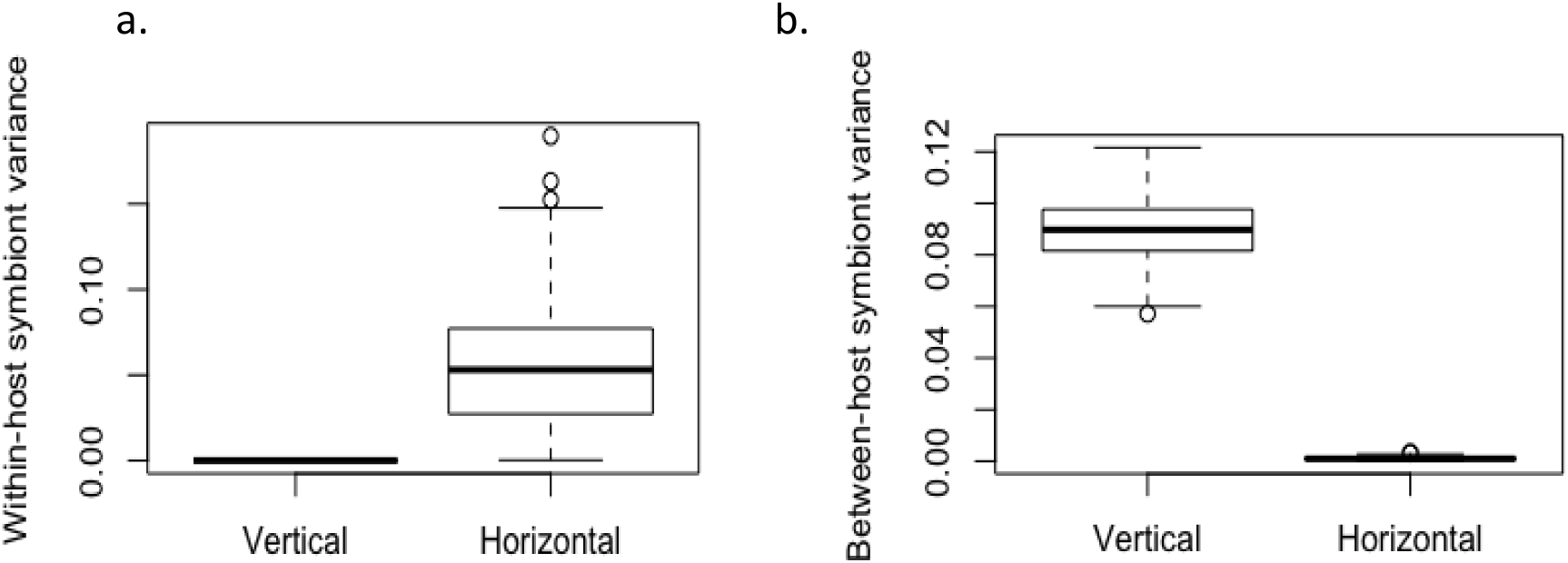
Variance of symbiont fitness scores (a) within hosts (average of the variance of scores for symbionts within single hosts), (b) between hosts (the variance of the average scores of hosts). Each point is the variance in a single random transmission simulation with only vertical or horizontal transmission types after 1000 time steps.

**Figure 6:**
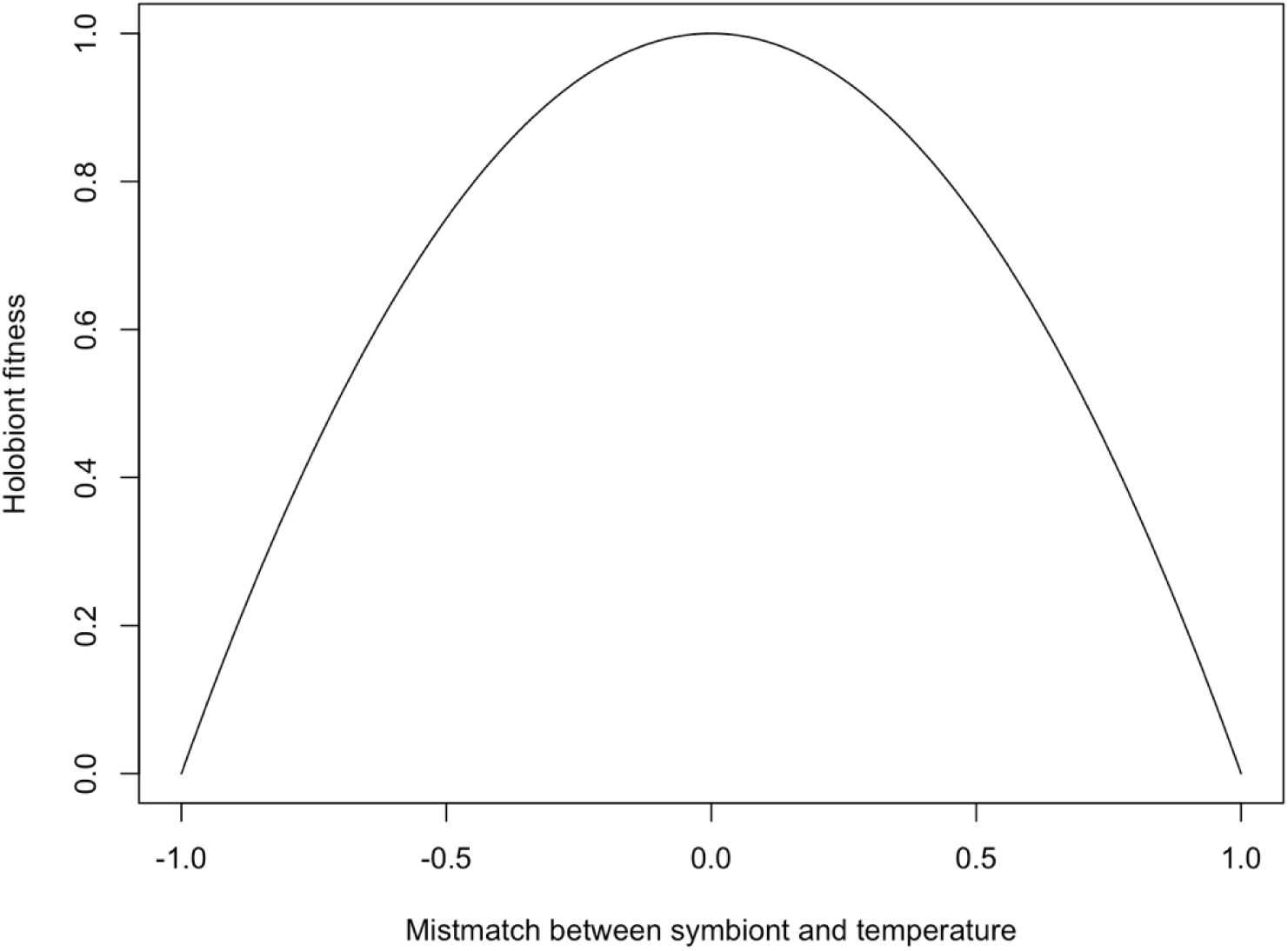
Fitness curve. The fitness impact of each symbiont on the holobiont (fitness score) was calculated based the difference between the symbiont gene value (optimal temperature) and the current temperature. When the symbiont gene value matched the current temperature, the fitness score was set equal to 1 and it declined (beta function) as the gene value and temperature were more different. A positive mismatch value indicates the environment is colder than the optimal value for the symbiont and a negative value indicates the environment is colder than the optimal value.

## DISCUSSION

We present here an agent-based model that investigates the adaptation ability of vertically transmitted holobionts and horizontally transmitted holobionts to increasing stress through symbiont adaptation. By introducing intraspecific trait variation and enabling evolution, we were able to study how selection-mutation-drift affected the population dynamics of holobionts with different transmission modes. Our model clearly demonstrates that both Vs and Hs are able to adapt to increasing environmental stress via symbiont adaptation, and their relative competitiveness depends on whether symbiont transmission is random or not. As long as the stress does not increase at a speed that will drive the holobionts to extinction rapidly, symbiont adaptation alone could increase holobiont stress tolerance.

Our model suggests the answer to a long debated question: can horizontally transmitted species be considered as a selection unit even though the symbiont disassociates from the host during holobiont life history (16, 47)? We showed that although offspring do not necessarily inherit symbionts from their parents directly, they could still acquire adapted symbionts. We found stress tolerance conferred by symbionts improved over time as the mean symbiont gene value increased along with increasing temperature. This was because holobionts whose symbiont values better matched the temperature were more likely to survive, and they passed their adapted symbionts to individuals in the next generation. In other words, because selection acts on the population level and drives the symbiont to confer higher stress tolerance, individual holobionts sampled from a symbiont pool that was adapting, and thus became adapted. This process looks like a Lamarckian evolution because agents acquire stress tolerance by associating with stress-resistant symbionts horizontally (48), but it is actually Darwinian evolution on the whole symbiont population level since the symbiont gene frequency changes due to selection against maladapted holobionts. Osmanovic *et al*. (49) found that selection on toxin tolerance of vertically transmitted bacteria alleviated holobiont stress in a long-lasting toxic environment. They suggested that horizontally transmitted symbionts can also be selected for and will confer higher stress tolerance to hosts over time which is similar to our results here. Acquiring adaptive traits via associating with horizontally transmitted symbionts is commonly observed in nature. In aphids, the secondary symbionts are considered to form a horizontal gene pool which facilitate aphid adaptation (50). Redman *et al*. discovered fungal-free plants that received *Curvularia* isolates (a type of fungal endophyte) from geothermal regions exhibited improved thermal tolerance, which also suggests stress tolerance could be acquired horizontally (51). In addition, research on corals with horizontally transmitted symbionts revealed surprisingly high fidelity in symbionts across generations, indicating unknown mechanisms that maintain symbiont stability in Hs (52). Thus we propose that symbiont-mediated holobiont adaptation is plausible in both Hs and Vs.

Further comparison of Vs’ and Hs’ adaptation ability shows that whether holobionts are able to identify and partner with suitable symbionts or not is critical in determining their relative competitiveness. Researchers have opposite opinions regarding which transmission mode is more advantageous because of following reasons: on one hand, Hs could exploit a broader range of symbionts, and are likely to take up adaptive symbionts from the environment (21, 53), while Vs can only inherit parents’ symbionts, which may be maladaptive once the environment changes (54, 55). On the other hand, adaptive symbionts could be passed down faithfully in vertical transmission, but get lost during horizontal transmission (4, 15). Our results suggest both arguments are partially correct, but are case dependent because symbiont acquisition could be random or non-random, which has significant impacts on the outcome (56). When transmission is random, Hs have no advantage even though the holobionts have access to a wider range of symbionts, because the chances of taking up adaptive and maladaptive symbionts are equal; but when they are able to recognize adaptive symbionts, they can pick up the fittest symbionts from a larger pool compared to the limited choices Vs have, which greatly improves their performance.

In random transmission scenarios, drift is stronger in Vs and can drive symbiont fixation in offspring lineages rapidly because they experience strong bottleneck effects given limited symbiont options in parents (18, 57); in contrast, Hs offspring tend to maintain their parents’ optimum according to the law of large numbers, because they sample from a larger symbiont pool (58). Our simulation of Hs and Vs without selection has shown that symbiont similarity is higher within each V than within each H (*i.e*., within host symbiont variation is larger in H individuals than in V individuals), but lower among Vs than among Hs (*i.e*., between-host symbiont variation is lower among Hs than among Vs). This is consistent with studies on corals, that found symbionts among vertically transmitted corals are more differentiated, and symbionts among horizontally transmitted corals are less differentiated (52, 59). In psyllids, some hosts are found to harbor only clonal symbiont lineages due to their vertical transmission mode, and the low diversity within the holobiont suggests a strong bottleneck effect (60). Given enough time, vertically transmitted symbionts could become so specific in host lineages that they will have reduced genome sizes and cannot escape from hosts (61). The strategies of Vs and Hs are thus different when facing environmental changes. Vs produce lineages that are distinctive in their stress tolerance, while Hs produce individuals more likely to persist in all conditions. Because maladaptive lineages that are dominated by maladaptive symbionts can be removed quickly from V populations, while maladaptive symbionts can persist in H populations through coexisting with adaptive symbionts, Vs are able to respond to environmental changes faster than Hs, and thus adapt faster when symbiont transmission is random.

Moreover, extremely maladaptive symbionts could be transmitted across the whole H population, while they are constrained in certain lineages in the V population, because of the larger potential host population available for each symbiont in Hs (21). From this perspective, having maladaptive symbionts is similar to having parasites, which may provide additional explanation for why higher virulence is often associated with horizontal transmission (24, 27). Because maladaptive symbionts can be acquired by holobionts other than direct descendants, they could persist in Hs just like infectious diseases, and decrease Hs’ overall fitness. In contrast, V individuals have a smaller sampling pool, which is determined by their parent, and offspring that are quickly dominated by maladaptive strains due to drift can be selected against (57), so overall fitness improves faster under selection. In short, once maladaptive symbionts arise in Hs, they can be taken up and spread through the population and are unlikely to be eliminated, but in Vs they can be trapped within lineages and be selected against, causing lower overall fitness in Hs than Vs. In an extreme scenario when the stress increasing speed is zero *(i.e*., under a constant environment), although both populations could persist for the whole simulation, Vs had slightly higher fitness especially when the mutational variance is large, where maladaptive symbionts are likely to arise and spread in Hs. Given enough time, Vs should be able to outcompete Hs (Fig. 5a). This resembles results of a study on virulence in pathogens, where vertical transmitted pathogens were selected for to prevent host from acquiring more virulent pathogens horizontally (27), and is consistent with study results of cultural transmission that suggest Vs are favored under stable conditions (33, 62). Our simulation also support the hypothesis that vertically transmitted holobionts such as corals may be more robust to climate change (63), but conditional on them conducting random transmission.

When we incorporated optimal symbiont uptake into the model, surprisingly we had the opposite result that Hs were better at tracking a changing environment. Such “recognition” of suitable symbionts is not well explored in previous studies (28, 34), yet their impacts on the outcome was very important in our simulation. We think such optimal transmission is equivalent to the positive transgenerational feedback proposed by Xue and Leibler (64). They suggested phenotypes of parents are reinforced in offspring, so organisms could adapt to varying environments. The positive transgenerational effect has been observed in some empirical studies, which is probably caused by epigenetic processes (65, 66). Our previous experiment on green hydra also showed that positive transgenerational effects could occur in holobionts simply via transmission of acclimated symbionts. The model presented here is slightly different from that described in Xue and Leibler’s and examples mentioned above, because here we are focusing on symbiont evolution rather than host evolution, and the phenotypes are distributed continuously rather than discretely. Nevertheless, the acquired symbionts were derived from those hosted by holobionts in the previous generations, which were more likely to be decedents of adaptive symbionts than maladaptive symbionts in their ancestors. Because the temperature in our model changes in one direction, the fittest new mutants are more likely to arise from the dominant, fittest strain in parent, instead of from a less fit strain. Such horizontal transmission of stress-tolerant symbionts has been observed in corals, which suggests holobiont adaptation can be achieved through transmission of adaptive symbionts (67, 68). By having access to symbionts released by holobionts other than their direct ancestors, Hs were able to selectively associate with the fittest strain that arose in the population which is in contrast to Vs. The ability to pick up the fittest symbionts, which is similar to copying from the most successful individuals in social learning, increases Hs fitness and allows them to adapt faster than Vs (69).

There are two potential mechanisms that hosts are able employ to take in the fittest symbionts: the first is via partner choice; the second is via passive up take. Mounting evidence of partner choice is being shown for holobionts (70), for example, plants allocate more resources to more beneficial partners and punish more parasitic partners (38, 71), so they could avoid cheaters and maximize their fitness during symbiosis. Bobtail squid are able to prevent colonization of deficient luminescent bacteria *Vibrio fischeri* in their light organ, which could be another case of partner choice (72). Massive coral bleaching could also be a case of hosts ejecting maladaptive symbionts and establishing symbiosis with adaptive ones (40). Corals are able to shift their symbiotic algae from sensitive strains to stress tolerant strains, so they can become more robust to stress such as high temperature and UV (53, 73–75). Thus even coral hosts that have long generation times and may not be able to adapt to warming temperatures quickly enough still have a chance to withstand climate change (3). An interesting example of wasps also shows that parents rather than the offspring may be able to choose symbionts to be transmitted (37), as such transmission could be controlled on a cellular level (76). In addition, even if the host is not able to identify the appropriate partner, they could form symbiosis with symbionts that pass an environment filter. Because symbionts vary in their stress tolerance, a strain that exhibits the highest fitness under certain conditions could outcompete other strains, and then form symbiosis with hosts (77). Hosts taking up these adaptive symbionts would acquire appropriate stress tolerance towards their environment (12). In insects, thermal stress could easily wipe out their thermal sensitive symbiotic bacteria, which could reduce their fecundity (8, 78). However, there are strains that could withstand such stress, and introducing these strains into hosts can restore hosts’ reproduction and enhance their thermal tolerance (8, 79). Symbionts that diverge in their niches are able to confer different traits and form association with hosts in distinctive conditions, which facilitates holobiont diversification (9, 50, 80). These examples provide solid evidence of optimal transmission of symbionts and suggest we should take optimal symbiont transmission into consideration when studying holobiont persistence in stressful environments.

Regardless of transmission mode and the ability to associate with optimal symbionts or not, holobionts were more likely to persist under stress when their mutational variance was large and stress increasing speed was low. It is not surprising that holobionts were able to persist longer at low rates of increasing temperature, because other studies also suggest rapid increase in stress could quickly wipe out the organisms (45, 81). However, Ayllón *et al*. (46) did not detect significant impacts of mutational variance on adaptation, and they suggested it was because their simulation ran for a limited number of generations (barely over 100 generations). In contrast, our simulations lasted for 4000 generations. Moreover, whether holobionts went extinct or not seemed to be confined by non-linear combinations of the mutational variance and temperature increasing speed, which requires larger increase in mutational variance at high rates of temperature increase than that at low rates of increase (Fig. 4).

Our models are the first exploring holobiont adaptation to stress, and demonstrate that both Hs and Vs are able to adapt, even when symbionts and hosts are disassociated between generations in Hs. Yet the model could be further expanded by incorporating more factors that might affect the dynamics between Hs and Vs in different scenarios. For instance, conducting horizontal transmission could be virulent, which might reduce Hs’ overall fitness and their competitiveness (27). In jellyfish, horizontal transmission favored symbionts shift to being parasitic, which reduced the growth rate of jellyfish up to 50% (22). We have applied this only to scenarios when there is no change in temperature, and found a 5% decrease in Hs’ growth rate could greatly reduce their relative frequency in both random transmission and optimal transmission. This implies that under constant environments, vertical transmission with less virulent symbionts would be favored (17, 33, 62, 82). But if we adjust the virulence parameter in temperature increasing scenarios, we will be able to explore how virulence could alter holobiont population dynamics. Another factor worth exploring could be the tradeoff between thermal tolerance and fitness cost. Studies on corals reveal that symbionts that confer higher thermal tolerance may slow down the growth of corals (83, 84). Imposing such constraints may enhance the authenticity of the model because holobionts will not be able to increase their thermal tolerance infinitely. We can also investigate into how Hs and Vs react in other scenarios such as random environments or fluctuating environments. We hope our models could provide a basic framework for holobiont evolution and shed light on symbionts’ roles from both ecological and evolutionary perspectives.

## MATERIALS AND METHODS

Our model was coded in python 3.0 with the agent-based modeling package “Mesa”. Here we present a model description following the ODD (Overview, Design Concepts, Details) protocol by giving a summary of the overall model structure and processes (29). Detailed variables and functions definitions can be found in Appendix A.

### Purpose

The purpose of the model is to understand whether holobionts with vertical transmission or horizontal transmission could adapt to changing environments solely depending on their symbionts, and how drift and mutation would affect this process. The stress in this model is called “temperature”, which is a hypothetical stress without a real world metric because it could also be called “salinity” or “UV intensity”. Similarly, each timestep represents one generation time, and has no corresponding real-world time length, because symbiotic species vary in their generation time. To minimize the model complexity, holobionts (the agents) in our model are autotrophic or heterotrophic with unlimited food and so do not need to compete for food, but they have to compete for space, which is common in ecosystems such as coral reefs or for plants (41, 85). Once a grid cell is occupied, it cannot be taken over by another agent unless the previous occupier is dead.

### Scales and variables

The model simulates adaptation of Vs and Hs under different temperature regimes, which contains three hierarchical levels: 1) environment, 2) holobiont, 3) symbiont. The object of the environment level is temperature. The temperature is a standardized hypothetical parameter without units because we are not studying a specific species and a specific stress tolerance, but rather to verify if holobionts can evolve as a unit. Each time step represents one generation, and agents could die or give birth to one offspring. The objects of the second level are the two types of holobionts, Vs and Hs, which only differ in their transmission modes. To track the adaptability of Vs and Hs, agents reproduce asexually and do not switch their transmission modes across generations. This allowed us to better track symbiont compositions of the two agent types over time, and compare fitness of Vs and Hs. The object of the third level, symbiont, is the only factor that determines holobionts’ fitness. Because our goal is to test whether both Vs and Hs can adapt via symbiont adaptation, and many holobionts’ stress tolerance is largely determined by symbiont stress tolerance (40, 86–88), we assume no host impacts on holobiont’s stress tolerance in this model. Each agent contains a fixed number of symbionts, and each symbiont is assigned a randomly generated gene value from which the host fitness will be calculated.

### Scheduling

The model is executed in following steps:

#### (1). Model Initialization

Initially 2000 agents are generated and randomly distributed on a 50*50 size grid. In cases where both Hs and Vs are presented, each agent has 50% chance to be H and 50% chance to be V. Each agent contains 50 symbionts, whose value is drawn from a normal distribution with mean equal to 0.5 and variance equal to mutational variance, truncated at two standard deviations (46, 89). Initial temperature is set at 0.5 as well.

#### (2). Holobiont reproduction

At every time step, each agent has a chance to reproduce based on its fitness, which is the mean of the fitness score calculated for its symbionts, assuming all the symbionts within the host will affect the holobiont’s fitness. We used a symmetric beta distribution to calculate the fitness score (90), which reaches the maximum when the symbiont’s gene value matches current temperature (Fig.6), and reaches a minimum when the mismatch is equal or greater than one, in other words, the current temperature could be either too hot or too cold for the symbiont. A random number is generated and compared to the holobiont’s fitness value to decide whether it will reproduce or not. Because conducting horizontal transmission might suppress holobiont reproduction, we introduce a cost parameter “l” that could reduce H’s reproduction probability (82).

#### (3). Symbiont transmission

In the default model, drift is introduced by enabling random symbiont transmission in Hs and Vs. V offspring sample symbionts from their parents randomly with replacement, so their symbionts could be a subset of those in their parents. H offspring sample symbionts from other agents around them once they are produced and dispersed (17). This transmission process is equivalent to picking up symbionts from a common pool constitute of symbionts released from agents (28).

In the optimal transmission model, drift is suppressed by enabling agents to pick the fittest symbionts from parents. As in cases where certain symbiont strains is dominant while the rest strains are marginal, here offspring choose the fittest strain from the parent (in V) or from neighboring agents (in H) to make up the symbiont population, and then randomly sample with replacement as in the default model to fill the rest of its symbiont population.

#### (4). Symbiont mutation

During holobiont reproduction, the symbiont may mutate by chance once it is acquired by the offspring. The mutation probability of each symbiont strain within the holobiont is set at 0.01 (91), given that symbiont generation time is shorter than holobiont generation time while the abundance is large (92, 93). The mutated symbiont gene value is drawn from a normal distribution with mean centered on the original symbiont gene value, variance controlled by the mutational variance, truncated at two standard deviation from the mean (46). Since we are interested in whether symbiont gene changes could drive holobiont adaptation, we assume heritability equals one and there is no environmental variance in our model.

#### (5). Holobiont dispersal

We use a stepwise function to control the position of a newborn holobiont. Because the further away from the parent the less likely the offspring would occur (94), we set equally high probability within certain distance for the new holobiont to occupy the gird, and equally low probability beyond certain distance for the holobiont to occur.

#### (6). Selection

Agent fitness is calculated as described above, except for the decision whether the agent will die or not.

Detailed parameters are documented in appendix A, and each parameter combination was run for 4000 time steps and at least 20 iterations.

### Design Concepts

#### Emergence

Population dynamics emerge from the reproduction, selection and competition of the two types of agents, embedded with symbiont drift and mutation. Adaptation is driven by interactions between each agent and the environment.

#### Sensing

Agents can perceive the temperature and know their transmission modes. In the optimal transmission model, agents are able to pick the fittest symbionts from their ancestors.

#### Interaction

Competition for space (empty cells) is modeled explicitly, based on a first come first serve rule. Reproduction and selection is simulated based on the beta functions (90) which depict the relation between fitness and temperature stress.

#### Stochasticity

At each process, the order of agents to execute the code is random. All behavioral parameters are generated based on empirical probability distributions. That means the reproduction, dispersal, and selection of agents are not deterministic, and symbiont values after mutation and inheritance in each agent are random. This introduces stochasticity on both symbiont and holobiont levels, and the final results are emergent from interactions between individuals and the environment.

#### Observation

For model analysis, population-level data were recorded, i.e., population size over time, relative ratio of the two agent types, mean holobiont optimal temperature, and time to extinction, etc. (95).

### Simulation scenarios

#### (1) Test case

To test and validate the model, we constructed a basic model that both agent types have equal fixed reproduction and survival probability, so they are selectively neutral and their fitness does not depend on the environment. This could be viewed as two genotypes in a finite population, whose fixation should be random. According to Kimura & Ohta (57), when the initial relative ratio is 0.5, either H or V will fix at 50% chance, and if the initial relative ratio moves away from 0.5, one or the other will be more likely to fix within dramatically less time.

#### (2) Increasing temperature

The temperature was projected to increase at different speed infinitely, which means holobionts would only be able to persist by having novel mutations conferring thermal tolerance higher than that of existing symbionts. This model tested whether both Vs and Hs could keep up with increasing stress, and explored how symbiont mutational variance might affect this process given different increasing speed. For both random transmission model and optimal transmission model, Hs and Vs were simulated in coexistence and in separation.

## Statistical analysis

We used paired Wilcoxon rank tests to compare the fitness between Hs and Vs across all the parameter combinations. We summed up the mean holobiont fitness at each time step of every iteration for a given parameter combination and performed a log transformation. This provides us information that includes both the fitness at each time step as well as the extinction time. We used the same test for Hs and Vs extinction time, correlation between holobiont gene values and the temperature, fitness score variance within holobionts, and fitness score variance between holobionts.

## ACKNOWLEDGEMENTS

We thank Dr. Rudy Guerra and Dr. Marek Kimmel for providing assistance to our model development.

